# Macrophage aggresome-like induced structures are flexible organizing platforms for immune signaling

**DOI:** 10.1101/2020.11.30.398750

**Authors:** Marie-Eve Charbonneau, Vedhika Raghunathan, Mary X.D. O’Riordan

## Abstract

Macrophages adopt a pro-inflammatory phenotype in response to environmental challenges in a process that often coincides with the formation of transient cytosolic p62/SQSTM1 inclusions containing ubiquitinated proteins in structures known as aggresome-like induced structures (ALIS). Although described as stress-induced inclusions that accumulate aggregated proteins, little direct evidence supports their hypothesized structural role in the context of immune stimulation. Here, we showed that these structures in primary macrophages are induced by multiple microbial-based ligands, including exposure to cytosolic double-stranded DNA. Rather than accumulating aggregated proteins, we observed that ubiquitinated proteins form a ring-shaped structure around the perimeter of these circular foci. We identified that different microbial stimuli induced the formation of ubiquitin-positive foci with distinct characteristics and we observed selective recruitment of context-dependent immune regulators. Our findings are consistent with a model where these ubiquitin-containing structures act as adaptable organizing centers for innate immune signaling.

**SUMMARY:** Charbonneau et al. demonstrate that ubiquitin- and p62-containing cytosolic ring-shaped structures induced by bacterial infections, microbial ligands and cytosolic double-stranded DNA contain context-dependent immune regulators, revealing an important insight on the cellular architecture required to coordinate signal transduction in macrophage.

---

Macrophages are essential for control of homeostasis, tissue repair, and inflammation and are a part of the front-line defense against invading pathogens (Wynn et al., 2013, Murray and Wynn, 2011). Macrophages recognize microbial challenge through a variety of pattern-recognition receptors (PPRs), including Toll-like receptors (TLRs), nucleotide-binding domain, leucine-rich repeat-containing proteins (NLRs), the AIM2-like receptors (ALRs), and several DNA sensors such as cGAS, the DNA-sensing enzyme cyclic guanosine monophosphate-adenosine monophosphate synthase (Fitzgerald and Kagan, 2020, Kieser and Kagan, 2017). Although these immune receptors share no structural similarities, a unifying principle governs downstream signaling events. Indeed, activation of these receptors leads to the formation of large oligomeric complexes that allow signal transduction and generation of inflammatory mediators such as cytokines and cell-intrinsic defense mechanisms, including the expression of interferon (INF)-stimulated genes (Kagan et al., 2014, Fitzgerald and Kagan, 2020). The formation of these supramolecular organizing centers (SMOCs) involves the oligomerization of adaptor proteins and the recruitment of specific effector proteins, which leads to the formation of localized punctate structures in the cells. Examples of SMOCs include the Myddosome, which consist of a TLR sensor, the downstream adaptors MyD88 and TIRAP as well as protein kinases of the IRAK family, and the inflammasome, which includes an NLR protein, the ASC adaptor, and a member of the caspase protein family (Kagan et al., 2014). The concentration of signaling components into specific subcellular structures is thought to enable an effective immune response by increasing the threshold for effector protein activation and signal amplification (Fitzgerald and Kagan, 2020). Despite these recent advances in our knowledge of the cell response to microbial sensing, more work is needed to better understand the spatiotemporal organization of signaling molecules that link the initial SMOCs formation and the downstream long-lasting cellular responses, including changes in transcription, translation, and metabolism.

A type of punctate cytosolic structure that was described in immune cells upon stimulation with TLR ligands or during infection with bacterial pathogens such as *Listeria monocytogenes* and *Mycobacterium tuberculosis* is the aggresome-like induced structure (ALIS) (Szeto et al., 2006, Liu et al., 2012, Fujita et al., 2011, Canadien et al., 2005). These structures are solely characterized by the presence of ubiquitinated proteins and the multi-functional adaptor protein p62/SQSTM1. Similar structures were observed in many cell types in response to proteotoxic stresses including treatment with an endoplasmic reticulum (ER) stress inducer, a protein translation inhibitor, and exposure to reactive oxygen species (ROS) (Vasconcellos et al., 2016, Liu et al., 2012, Jena et al., 2018). Accordingly, ALIS are described as structures that accumulate misfolded ubiquitinated proteins upon cellular stress, when the degradative pathways such as the proteasome and the autophagy machinery are overwhelmed. In this model, p62 serves as a scaffolding component required to package these aggregated ubiquitinated proteins. Of note, ALIS puncta are distinct from aggresomes as their formation is independent of the microtubule network and they are not surrounded by a vimentin cage (Szeto et al., 2006). Although first described in dendritic cells upon inflammatory stimulation (Lelouard et al., 2002), the information available regarding these structures was mostly acquired by studying diverse cell types exposed to a variety of cell stressors that are not related to immune responses. Therefore, a detailed analysis of the structure and components present in these cytosolic puncta, specifically in the context of innate immune signaling, remains to be determined.

Multiple lines of evidence suggest that ALIS might be more than depots of aggregated proteins in immune cells. First, these foci form extensively after TLR stimulation, which would suggest that triggering an immune response in macrophages is concomitant with the accumulation of a large number of misfolded proteins that need to be contained into specific depots for future degradation. However, little evidence supports that TLR signaling induces accumulation of misfolded proteins or that prevention of ALIS formation following immune stimulation correlates with cellular toxicity in macrophages. Second, p62 is an important protein for homeostatic cell functions besides its role in packaging aggregated proteins (Moscat et al., 2016, Sanchez-Martin et al., 2019). This multivalent adaptor protein can self-assemble and form liquid-like bodies or droplets, a process enhanced by binding to ubiquitin chains, which is consistent with the co-occurrence of these two proteins in cytosolic condensates (Sun et al., 2018). Multiple reports have shown that p62 droplets can act as signaling hubs for protein activation, including major components of signaling pathways like the nuclear factor erythroid 2-related factor 2 (Nrf2), the mechanistic target of rapamycin complex 1 (mTORC1), the nuclear factor kappa-light-chain-enhancer of activated B cells (NF-kB) and to promote caspase-8 activation (Jin et al., 2009, Park et al., 2013, Sanchez-Martin et al., 2020, Duran et al., 2011, Kehl et al., 2019). Together, these studies highlight the potential for p62 containing structures to organize signaling pathways instead of promoting protein degradation.

Lastly, multiple bacterial pathogens prevent the formation of punctate structures containing ubiquitinated proteins during infection. Indeed, the *Salmonella enterica* serovar Typhimurium type III secretion system effector SseL, a bacterial deubiquitinase, prevents the formation of ubiquitin-positive cytosolic puncta during infection. (Mesquita et al., 2012). Similarly, in a type IV secretion system-dependent manner, *Legionella pneumophila* and *Brucella abortus* actively prevent the formation of ALIS, although the specific effector proteins involved in this process remain to be identified (Ivanov and Roy, 2009, Salcedo et al., 2008). Taken together, these observations open up new possibilities regarding the nature of ALIS foci and their potential implication in macrophage immune defenses.

Here, we show that the formation of ubiquitin-containing cytosolic structures occurs in macrophage in response to various immune stimuli, which include TLR stimulation, bacterial infection, and exposure to cytosolic double-stranded DNA (dsDNA). Furthermore, we show that while ubiquitinated proteins and ubiquitin-binding proteins like p62 are shared elements of these structures, other components are context-specific and depend on the signal triggering the immune response. We also provide evidence that these structures selectively recruit key components of signaling pathways, including proteins involved in the NF-kB pathway or the type I interferon response. Therefore, our findings are consistent with the idea that these cytosolic ubiquitin-containing foci might be similar to the SMOCs and act as subcellular sites in macrophages where innate immune signaling occurs. A better understanding of these ubiquitin-containing structures from a cell biology angle provides an important insight into how macrophages perform signal transduction in a regulated manner to promote productive immune responses.

## RESULTS

### LPS stimulation induces the formation of ring-shaped cytosolic ubiquitin- and p62-positive structures in primary macrophages

Formation of ubiquitin (Ub)-enriched cytosolic structures occurs in response to multiple cellular stresses, including oxidative and proteotoxic stresses, leading to the suggestion that these foci are misfolded proteins waiting for degradation (Liu et al., 2012, Vasconcellos et al., 2016, Szeto et al., 2006, Jena et al., 2018). Similar structures are also observed following immune stimulation of dendritic cells and macrophages, but the nature of these Ub-containing foci in the context of immune stimulation remains elusive. To directly address this question, we treated primary bone marrow-derived macrophages (pBMDM) with 10 ng/ml of lipopolysaccharides (LPS) for 6h and stained them for p62 and polyubiquitinated proteins. Approximately 86% of cells stimulated with LPS displayed 2 or more foci, while the number of Ub-positive foci in these cells ranged from 2 to 20 with an approximate average of 6 foci per cell (Fig. 1A). These LPS-induced Ub-positive foci have an average size of around 0.94 μm^2^ (Fig. 1B). We observed similar results using concentrations of LPS ranging from 0.1 ng/ml up to 10 ng/ml (Fig. S1A), suggesting that a more physiological concentration of 0.1 ng/ml of LPS is enough to trigger Ub-containing foci formation in macrophage. The ubiquitin protein has seven lysine residues that can be linked to other ubiquitin molecules. Globally, K48-linked ubiquitin chains are associated with proteasomal degradation, whereas K63-linked chains are important for protein trafficking, stability, and many signaling processes, including innate immune signaling (Yau and Rape, 2016). By staining macrophages with antibodies specific for K48- and K63-polyubiquitin chains, we observed that these cytosolic puncta are enriched in both polyubiquitin linkages. These observations suggest that ubiquitinated proteins contained in these structures might not all be targets for degradation but might rather play a functional or structural role (Fig. S1B). LC3, a marker of the autophagosome that binds directly to p62, is recruited to the Ub-and p62-containing foci (Levine et al., 2011, Fujita et al., 2011). We confirmed the recruitment of LC3 to the Ub-containing foci in pBMDM (Fig. S1C); however, the formation of these structures was independent of the classic autophagy machinery. Indeed, at 6h post LPS stimulation, there was no significant difference in the number of foci per cell when Atg7^fl/fl^ LysM-Cre pBMDM were compared to Atg7^fl/fl^ (WT) cells (Fig. S1D). We could, however, observe more Ub-positive foci in the cytosol of Atg7^fl/fl^ LysM-Cre pBMDM compared to WT pBMDM at 24h post-stimulation with LPS, suggesting a possible role for the autophagy process in clearing these structures (Fig. S1E).

**Figure 1:**
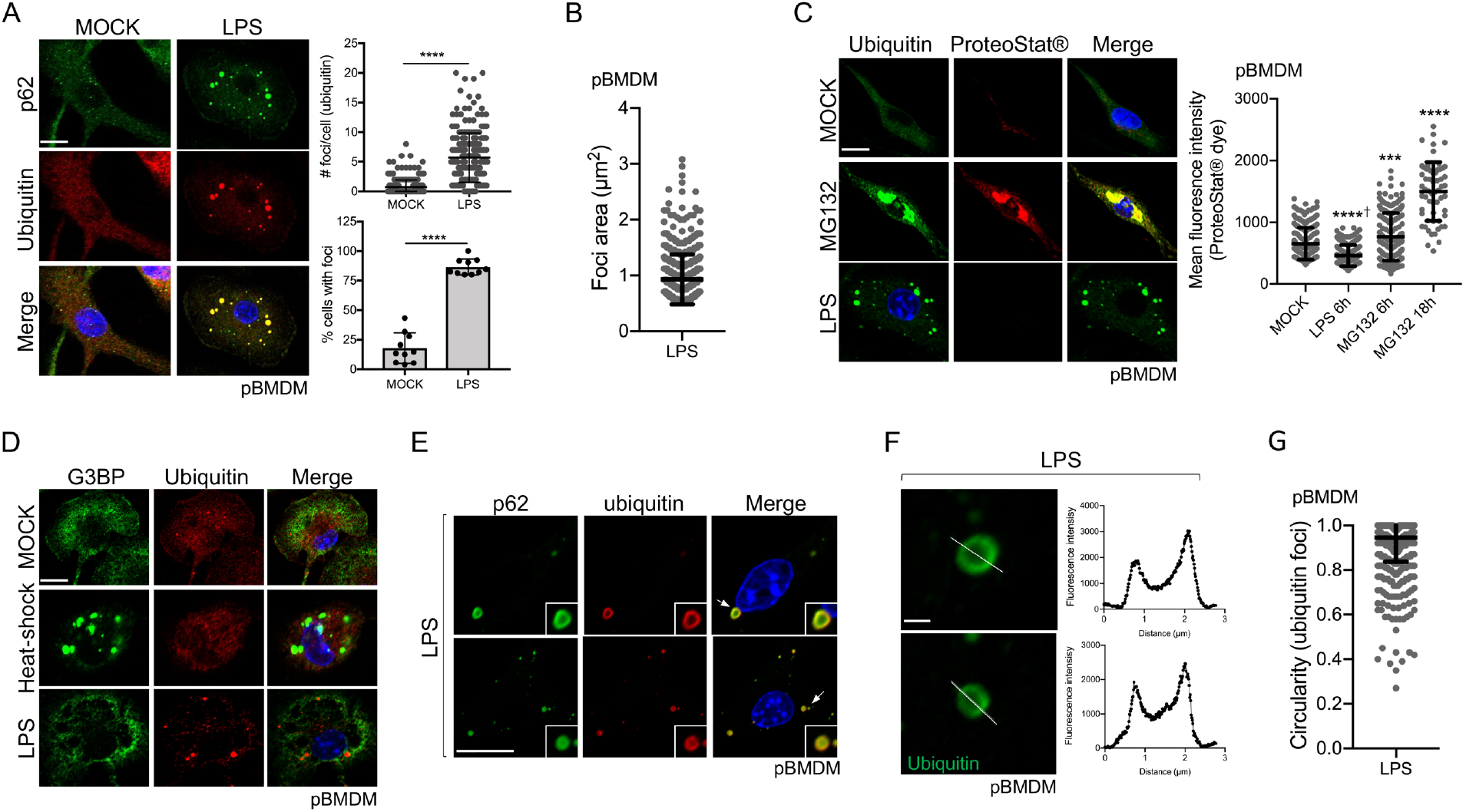
LPS stimulation induces the formation of cytosolic ring-shaped Ub- and p62-positive structures. ***A.*** Representative Confocal microscopy images of pBMDM left untreated or stimulated for 6h with 10 ng/ml of LPS. Macrophages were stained using antibodies against p62 and polyubiquitinated proteins. Quantification was done using the particle analysis function of FIJI to identify particles with a size between 0.5 and 15 μm^2^ and with a circularity higher than 0.3. At least 200 cells pooled from three independent experiments were used. ***B.*** Quantification of the foci area (μm^2^) from pBMDM treated with 10 ng/ml of LPS for 6h. ***C***. Representative images and quantification of pBMDM left untreated, stimulated for 6h with 10 ng/ml of LPS, or treated with the proteasome inhibitor MG132 for 6h or 18h and stained using an antibody against polyubiquitinated proteins and the aggresome/aggresome-like specific dye ProteoStat™. ***D***. Representative images of pBMDM left untreated, stimulated for 6h with 10 ng/ml of LPS or incubated at 42°C for 1h (heat-shock), and stained with antibodies against polyubiquitinated proteins and G3BP. ***E***. Images representing a magnified area of pBMDM treated and stained as described in *A*. The arrows indicate the region that is displayed in higher magnification. ***F***. Representative enlarged single confocal images showing the structures of a focus stained for ubiquitinated proteins after 6h treatment with LPS 10 ng/ml. The fluorescence intensity was calculated across the white line in the neighboring panel. ***G***. Quantification of the foci circularity (0 to 1) from pBMDM treated with 10 ng/ml of LPS for 6h. Graphs represent the mean and the corresponding standard deviation of the mean and significant differences were calculated using two-tailed Student’s t-test (*A*) or one-way ANOVA with Tukey’s post-test (*C*) (NS, not significant, ***p < 0.001, ****p < 0.0001, † significantly lower). Scale bar of 10 μm (*A*, *C*, *D, E)* or 1 μm *(F).*

ALIS were described as sites for accumulation of misfolded and aggregated proteins during stress, a role similar to that associated with aggresome formation (Szeto et al., 2006). In agreement, Ub-enriched foci observed during oxidative stress contain aggregated proteins (Jena et al., 2018). To directly test the assumption that ALIS induced by LPS treatment are depots of aggregated proteins, we used the reagent ProteoStat™, a widely used protein aggregate-specific molecular rotor dye that intercalates into quaternary structures usually found in misfolded and aggregated proteins and emits fluorescence (Bershtein et al., 2013, Seo et al., 2016, Shen et al., 2011). Primary BMDM treated with MG132, a proteasome inhibitor (Lee and Goldberg, 1998), for 6h or 18h clearly showed strong fluorescence that co-localized with ubiquitinated proteins (Fig. 1C). The mean fluorescence intensity of ProteoStat™ was significantly higher after proteasome inhibition when compared to untreated cells, confirming the utility of this reagent in assessing protein aggregates. Macrophages treated with LPS showed accumulation of Ub-positive puncta, but these structures were not labeled by ProteoStat™, suggesting that LPS stimulation is not associated with the accumulation of misfolded and aggregated proteins (Fig. 1C). Other types of cytosolic foci are observed in cells after proteotoxic stress, including stress granules, which are membrane-less compartments containing mainly untranslated mRNA and RNA-binding proteins (Protter and Parker, 2016). As stress granules shared similar properties compared to the Ub-containing structures observed in LPS-stimulated macrophages, we tested for the presence of a stress granule component, G3BP, in foci formed following immune stimulation. Co-staining of pBMDM for ubiquitinated proteins and G3BP showed that Ub-containing foci formed in response to LPS were distinct from the stress granules induced by heat-shock (Fig. 1D).

To better define the nature of the Ub-containing foci observed in response to LPS stimulation, we examined their structure by high-resolution confocal imaging. We observed that the Ub-and p62-containing foci commonly formed a ring-shaped structure (Fig. 1E-F). By analyzing the staining pattern of an individual puncta, we found that ubiquitinated proteins localized mainly on the edge of the spherical focus whereas p62, although enriched on the edge, also stained the region toward the center of the spherical structure (Fig. 1E). Detailed analysis of these LPS-induced Ub-positive foci showed a mean circularity (sphericity) of 0.94 (Fig. 1G). The consistent high circularity score of these foci might suggest that they behave similarly to membrane-less biomolecular condensates with liquid-liquid phase separation properties (Snead and Gladfelter, 2019, Wang and Zhang, 2019, Alberti et al., 2019). Moreover, the absence of staining for ubiquitinated proteins inside these foci is in good agreement with the idea that these structures are not just depots of aggregated proteins, in which case we would expect the entire foci to be labeled for proteins tagged with ubiquitin molecules. Taken together, our results unveil that these Ub-containing puncta observed in LPS-stimulated macrophages are organized structures, a characteristic reminiscent of other higher-order signaling complexes, like SMOCs, involved in immune signaling.

### LPS induces ALIS formation through a transcriptional program, with a minor contribution from the Nrf2 pathway

SMOC assembly occurs rapidly after recognition of a ligand, as exemplified by the formation of the myddosome within minutes after LPS stimulation (Fitzgerald and Kagan, 2020, Bonham et al., 2014). TLR dimerization allows the recruitment of the components already available and needed for formation of the signaling platform. ALIS assembly may occur through a distinct mechanism, as the earliest time we could observe Ub-positive structures in pBMDM was at 4h post-stimulation with LPS, suggesting a possible requirement for transcriptional events (Fig. S2A). In agreement with this hypothesis, the formation of Ub-containing foci in stimulated macrophages was blocked by treatment with actinomycin D, an inhibitor of mRNA synthesis, or with cycloheximide, an inhibitor of protein synthesis (Fig. S2B). Likewise, ALIS formation in response to LPS was abolished in TLR2/4/9^-/-^ immortalized bone marrow-derived macrophages (iBMDM) (Fig. S2C). Formation of Ub-containing foci in the context of oxidative stress, induced either by treatment with hydrogen peroxide or exposure to heme, is driven by the transcription factor NF-E2-related factor (Nrf2), a master regulator of the cellular response against oxidative stress (Szeto et al., 2006, Vasconcellos et al., 2016, Jena et al., 2018). However, ALIS formation induced by the autophagy inhibitor 3-MA is independent of the Nrf2 transcriptional response (Wenger et al., 2012), suggesting a disparity in the dependency for Nrf2 activation depending on the signal initiating the formation of these Ub-containing cytosolic structures. To further investigate the role of Nrf2 signaling for the formation of Ub-containing structures in our experimental model, we treated WT and Nrf2^-/-^ pBMDM with LPS for 6h and analyzed foci formation. Although the number of Ub-containing foci per cell was significantly lower, these structures could still be observed in approximately 74% of Nrf2^-/-^ macrophages (Fig. S2D). In agreement, 73% of cells treated with the reactive oxygen species scavenger *N*-acetylcysteine (NAC) had Ub-positive foci after LPS stimulation, although a lower number of foci per cell was detectable (Fig. S2E). Thus, our results suggest that activation of macrophages using LPS triggers the formation of Ub-containing cytosolic structures in a TLR- and transcription-dependent manner, and but does not require activation of the major cellular oxidative pathway.

### Distinct microbial ligands induce the formation of ubiquitin and p62-positive foci

We used LPS as an archetypal microbial TLR ligand to understand the detailed structure of the Ub-containing foci. Macrophages, however, recognize microbial challenge through a variety of pattern-recognition receptors beyond TLR4. To determine the relevance of ALIS formation after immune stimulation, we first treated cells with Pam3CSK4, a synthetic triacetylated lipopeptide, which is a potent TLR1/TLR2 heterodimer agonist that mimics bacterial lipoproteins found in both Gram-positive and Gram-negative bacteria. As with LPS, Pam3CSK4 treatment induced the formation of Ub-positive structures in a TLR2/4/9-dependent manner (Fig. 2A). We also observed that infection by two Gram-positive human pathogens, methicillin-resistant *Staphylococcus aureus* (MRSA) and *L. monocytogenes,* induced the formation of cytosolic structures containing ubiquitinated proteins (Fig. 2B-C). MRSA induction of ALIS was dependent on TLR signaling as foci formation was abolished in TLR2/4/9^-/-^ iBMDM (Fig. 2B). However, foci were observed in TLR2/4/9^-/-^ iBMDM infected with *L. monocytogenes,* although at a significantly lower number compared to WT cells (Fig. 2C). *L. monocytogenes* is an intracellular pathogen that replicates in the cytosol of infected cells and therefore can trigger the activation of multiple cytosolic innate immune pathways including the Type I interferon response (Woodward et al., 2010). DNA release from *L. monocytogenes* induces IFN-β secretion in a cGAS-, IFI16-, and STING-dependent manner (Hansen et al., 2014). Similarly, detection of the bacterial second messenger c-di-AMP secreted by *L. monocytogenes* promotes STING activation and the Type I Interferon response (Parvatiyar et al., 2012). As ALIS formation was observed independently of TLR2/4/9 in response to *L. monocytogenes* infection, we investigated the possibility that DNA sensing might be a trigger for the formation of Ub-positive cytosolic foci in macrophages. Indeed, transfection of G3-YSD, a palindromic DNA sequence that self-hybridized to form a Y-form short dsDNA and acts as a cGAS agonist (Herzner et al., 2015), as well as transfection of an immune-stimulatory DNA (ISD), resulted in the formation of Ub-positive foci in macrophages (Fig. 2D). A detailed analysis of the foci formed after infection with *L. monocytogenes* or transfection of dsDNA showed ring-shaped structures comparable to the foci observed after LPS stimulation (Fig. S3A-B). These Ub-positive foci could also be detected using a p62 antibody, highlighting again the similarity with the LPS-induced foci (Fig. S3C). Notably, we observed that Ub-positive structures often formed around the transfected DNA, as highlighted by DAPI staining (Fig. S3C). Although G3-YSD is a cGAS agonist, cGAS was not required for the formation of the Ub-containing foci. Indeed, cGAS^-/-^ pBMDM transfected with G3-YSD displayed a similar number of foci compared to WT pBMDM (Fig. S4A-B). Similarly, the downstream signaling adaptor molecule STING was not required for foci formation in response to dsDNA (Fig. S4C-D), suggesting that recognition of cytosolic dsDNA is a signal that triggers the formation of ubiquitinated protein-containing foci independently of the cGAS/STING signaling pathway. Taken together, our results are consistent with the idea that the formation of cytosolic ubiquitin and p62-positive structures is a common response to immune stimulation in macrophages.

**Figure 2:**
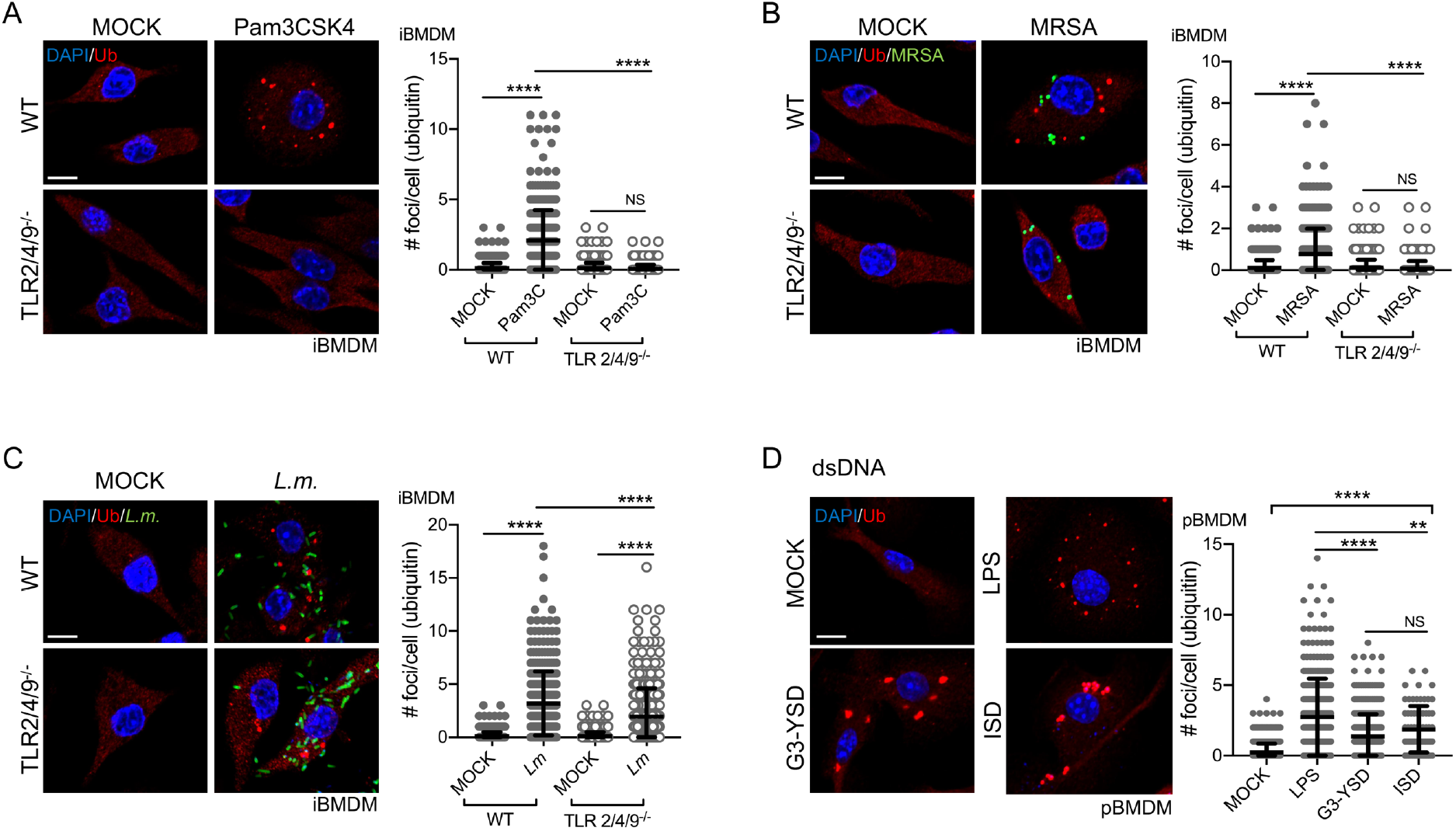
Distinct microbial-based immune stimuli induce the formation of Ub- and p62-positive foci in macrophages. ***A.*** WT and TLR2/4/9^-/-^ iBMDM were left untreated or stimulated for 6h with 100 ng/ml of Pam3CSK4. ***B***. WT and TLR2/4/9^-/-^ iBMDM infected with MRSA-GFP at an MOI of 20 for 1h, washed and incubated for 6h in a medium containing gentamycin to kill extracellular bacteria. ***C***. WT and TLR2/4/9^-/-^ iBMDM were infected for 0.5h with *L. monocytogenes* at an MOI of 5, washed, and incubated for 6h in a medium containing gentamycin. *L. monocytogenes* was stained using specific *Listeria* O antisera. ***D.*** pBMDM were stimulated with 10 ng/ml of LPS, transfected with 2 μg/ml of G3-YSD, or transfected with 25 pmol of ISD for 6h. Staining (blue: DAPI; red: polyubiquitinated proteins; green: bacteria) and image processing was done as described in Figure 1. Graphs represent the mean and the corresponding standard deviation of the mean from three independent experiments using at least 200 cells per condition. Significant differences were calculated using one-way ANOVA with Tukey’s post-test (NS, not significant, ** p < 0.01, **** p < 0.0001). Scale bar: 10 μm.

### Different microbial stimuli induce the formation of ubiquitin structures with different properties

We showed that multiple immune stimuli induced the formation of Ub- and p62-containing structures similar to those observed in LPS-treated cells. Treatment of cells with the TLR1/2 agonist Pam3CSK4 induced the formation of Ub-positive foci with a similar size range and abundance when compared to LPS stimulation (Fig. 2A and Fig. 3A), suggesting a common response downstream of TLRs engagement that triggers the assembly of foci with similar structural features. However, some differences in the Ub-positive foci structures were easily noticeable when other microbial ligands were used for stimulation. Indeed, macrophages transfected with dsDNA had a lower number of Ub-containing foci per cell compared to LPS stimulation (Fig. 2D). Moreover, the average sizes of the foci induced by either infection with *L. monocytogenes* (average size of 1.5 μm^2^) or dsDNA transfection (average size of 2.1 μm^2^) were significantly larger compared to LPS-induced puncta (average size of 0.94 μm^2^) (Fig. 3B-C). The circularity of the foci was also variable depending on the immune triggers, from a circularity value of 0.94 for LPS stimulation to 0.80 for *L. monocytogenes* infection and 0.82 for dsDNA transfection (Fig. 3D). Of note, the Ub-containing foci observed after *L. monocytogenes* infection or transfection with dsDNA were not labeled using the Proteostat™ dye, which is consistent with the idea that these foci are not representing an accumulation of aggregated proteins (Fig. 3E).

**Figure 3:**
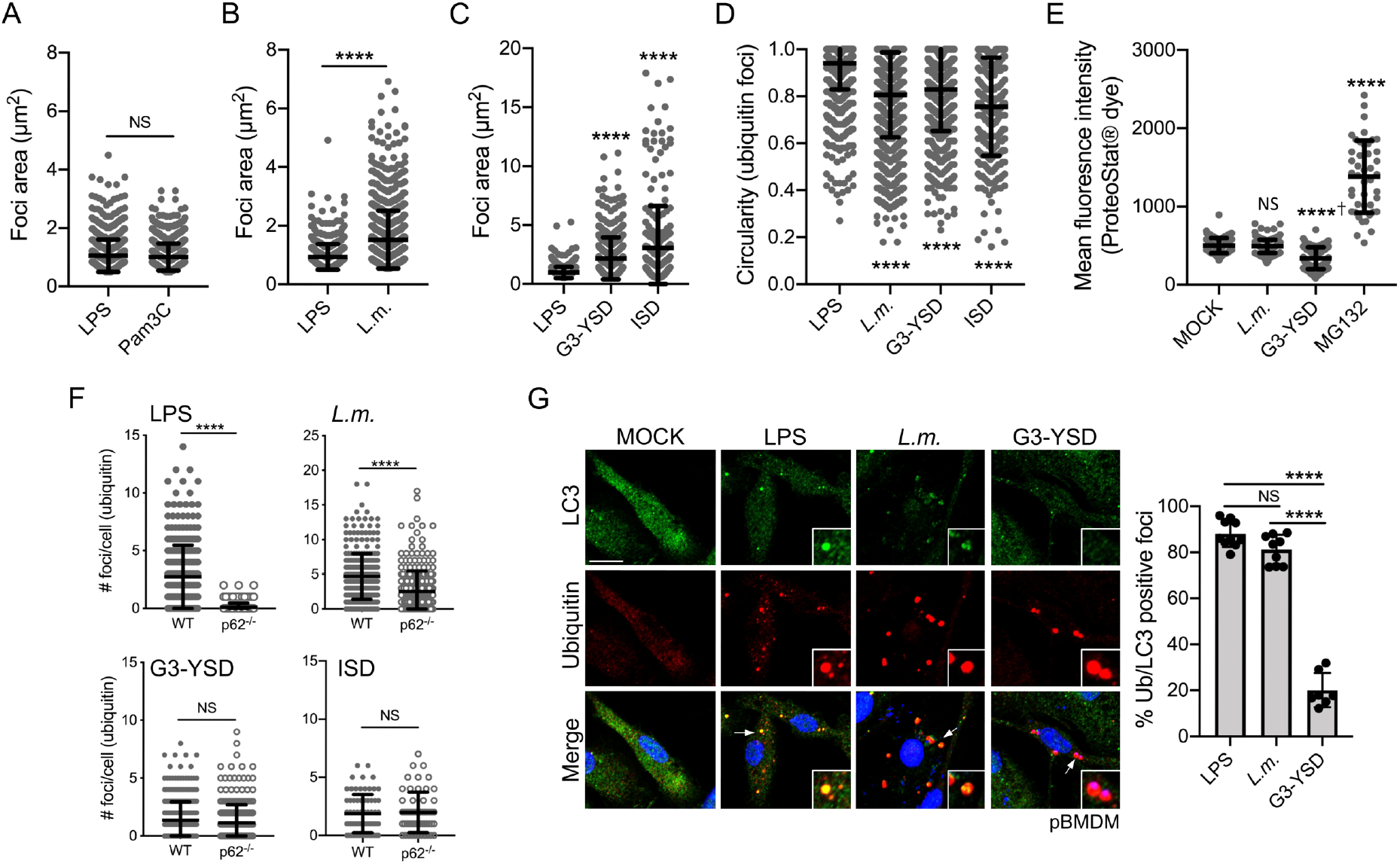
Different microbial stimuli trigger the formation of Ub-containing structures with different properties. Quantification of the foci area (μm^2^) using the staining for polyubiquitinated proteins from iBMDM treated with 10 ng/ml of LPS or 100 ng/ml of Pam3CSK4 for 6h (***A***), infected for 0.5h with *L. monocytogenes* at an MOI of 5, washed and incubated for 6h in medium containing gentamycin (***B***) or transfected with either 2 μg/ml of G3-YSD or 25 pmol of ISD for 6h (***C***). ***D***. Quantification of the foci circularity (from 0 to 1) using the staining for polyubiquitinated proteins from pBMDM treated with 10 ng/ml of LPS, transfected with 2 μg/ml of G3-YSD or infected with *L. monocytogenes* as described above. ***E***. Quantification of the mean fluorescence intensity for the aggresome/aggresome-like specific dye ProteoStat™ in pBMDM treated with the proteasome inhibitor MG132, infected with *L. monocytogenes* or transfected with 2 μg/ml of G3-YSD for 6h. ***F***. Quantification of the number of foci per cell containing polyubiquitinated proteins in WT and p62^-/-^ pBMDM treated with LPS 10 ng/ml, infected with *L. monocytogenes* as above or transfected with either 2 μg/ml of G3-YSD or 25 pmol of ISD. ***G***. Representative images of pBMDM treated with 10 ng/ml of LPS, infected with *L. monocytogenes,* or transfected with 2 μg/ml of G3-YSD for 6h and stained for LC3 and polyubiquitinated proteins. Graphs represent the mean and the corresponding standard deviation of the mean from at least two independent experiments using more than 145 cells per condition. Significant differences were calculated using two-tailed Student’s t-test (*A, B, F)* or one-way ANOVA with Dunnett’s *(D, E)* or Tukey’s (*G*) post-tests (NS, not significant, **** p < 0.0001, † significantly lower). Scale bar: 10μm.

As the structural features of these Ub-containing foci seem to be shaped by the stimuli used to induce their formation, we decided to look at the requirement for the major known component of ALIS, the adaptor protein p62. Using p62^-/-^ pBMDM, we found that this protein is required for the formation of Ub-containing foci in response to LPS stimulation (Fig. 3F), in agreement with previous reports (Fujita et al., 2011, Liu et al., 2012). However, p62^-/-^ pBMDM infected with *L. monocytogenes* displayed a significant number of Ub-containing foci per cell, although these structures assembled less efficiently when compared to WT pBMDM. Although p62 is observed in the Ub-containing foci formed following dsDNA transfection (Fig. S3C), their formation can occur independently of p62. Indeed, the formation of Ub-containing structures was similar in WT and p62^-/-^ pBMDM transfected with either G3-YSD or ISD (Fig. 3F). Another characteristic of the LPS-induced Ub-containing foci is the presence of the autophagy-associated protein LC3. To determine if this association is a unifying principle, we stained pBMDM infected with *L. monocytogenes* or transfected with dsDNA with an LC3 specific antibody. We observed that *L. monocytogenes* infection triggered the association of LC3 to the Ub-containing foci, whereas LC3 was absent from the structures generated after dsDNA exposure (Fig. 3G). Altogether, we showed here that although these Ub-containing foci that are induced by immune stimulation share some common features, the structural requirement and the composition of these condensates might be dependent on the input signal.

### Selective recruitment of innate immune regulators in ubiquitin-positive structures

Besides the presence of ubiquitinated proteins, p62, and in some contexts, LC3, nothing is known about the contents of these Ub-containing cytosolic structures. We have shown that various immune triggers can lead to the formation of structures containing ubiquitinated proteins with different characteristics and morphology. In this context, it is tempting to propose the idea that these structures might play a role in immune signal transduction that will differ depending on the stimuli. To address this possibility, we directly assessed the spatial distribution of some major immune regulators downstream of TLR4 and cGAS. TLR4 engagement by LPS triggers the assembly of the Myd88-containing complex the myddosome, which in turn promotes the activation of downstream protein kinases including the IKK complex and the MAP kinase (MAPK) p38 and their respective transcription factors NF-kB and AP1 (Fitzgerald and Kagan, 2020). A major modulator of the canonical NF-kB pathway is NEMO (IKKg), the regulatory subunit of the IKK activating complex (Maubach et al., 2017). pBMDM treated for 6h with LPS or transfected with the cGAS ligand G3-YSD were stained for ubiquitinated proteins and NEMO. We observed that approximately 31% of the Ub-containing foci were enriched for NEMO in response to LPS stimulation (Fig. 4A). Conversely, dsDNA transfection did not induce the relocation of NEMO to the Ub-positive structures, suggesting specificity in the components recruited to the Ub-containing foci depending on the immune trigger. Similarly, we investigated the localization of the MAPK p38 and found that the activated phosphorylated form of p38 could be detected in nearly 60% of Ub-positive structures after LPS stimulation (Fig. 4B). In contrast to what we observed for NEMO localization, exposure to dsDNA seemed to also be a signal for recruitment of phosphorylated p38, as approximately 44% of the Ub-positive foci induced by G3-YSD transfection were also enriched for this protein. Engagement of TLR4 can also induce signaling through the Toll/IL-1 receptor (TIR) domain-containing adaptor protein, TRIF (Yamamoto et al., 2003, Fitzgerald et al., 2003). TRIF signaling occurs through the TANK binding-kinase 1 enzyme (TBK1) and the transcription factor IRF3 to trigger Type I interferon expression. Besides its role in TLR signaling, TBK1 is an essential component of the cGAS/STING signaling pathway. Indeed, downstream of dsDNA sensing by cGAS, STING interacts directly with TBK1 to promote IRF3 activation and the Type I interferon response (Motwani et al., 2019). In resting macrophages, we observed that the activated and phosphorylated form of TBK1 is present at a low level (Fig. 4C). However, upon stimulation of pBMDM with either LPS or dsDNA, staining for p-TBK1 showed high prevalence for this protein in cytosolic puncta that also contained ubiquitinated proteins (Fig. 4C). Besides TBK1, cGAS is a major contributor to the cellular response to cytosolic dsDNA. As we observed the presence of DAPI-positive molecules inside the Ub-containing structures upon dsDNA transfection (Fig. S2C), we decided to look directly at cGAS spatial localization following exposure. The localization of cGAS in resting macrophages was mainly nuclear (Fig. 4D), in agreement with a previous report (Volkman et al., 2019). However, upon dsDNA transfection, a shift in the spatial distribution for the cGAS signal was visible, with increased detection in the cytosol, especially in proximity to the Ub-containing structures. Indeed, nearly 50% of the ubiquitin foci were enriched for cGAS following dsDNA transfection. In LPS-stimulated cells, cGAS was observed at a low frequency in the Ub-containing foci, which is quite reminiscent of the selective presence of NEMO in the foci induced by LPS treatment. Our results, therefore, suggest that multiple immune stimuli triggered the formation of Ub-containing cytosolic foci, but the components observed in these structures are dependent on the nature of the stimuli. These observations raise the interesting possibility that these Ub- and p62-containing foci might be structural platforms for innate immune signaling and the signal inducing their formation dictates the recruitment of specific components.

### Ligand recognition shapes the structural organization of the ubiquitin-containing foci

We have reported that cytosolic dsDNA induces the formation of Ub-containing foci that seems to encircle the cytosolic transfected dsDNA (Fig. S3C). This observation might suggest a correlation between the recognition of cytosolic dsDNA molecules and the formation of selective circular-shaped structures. The unpaired guanosine trimers at the ends of each strand of the dsDNA molecule G3-YSD are required for the activation of cGAS (Herzner et al., 2015). A derivative of this dsDNA molecule was previously described where the guanosine trimers were replaced by cytidine trimers to create the Y-shaped dsDNA C3-YSD. Although cGAS can bind to C3-YSD, this interaction results in a lower *in vitro* production of cGAMP and a level of INF-a secretion similar to untreated cells (Herzner et al., 2015). We confirmed that the transfection of primary macrophages with G3-YSD, but not with C3-YSD, induced the secretion of INF-β (Fig. 5A). Although the transfection of C3-YSD activated only weakly cGAS, this dsDNA molecule induced the formation of Ub-positive structures in the cytosol of macrophages (Fig. 5B). However, these structures are quite different from the ones observed following G3-YSD transfection (Fig. 5B). Indeed, C3-YSD transfection induced the formation of larger and irregularly shaped Ub-enriched structures, as shown by the area and circularity measurements (Fig. 5C-D). A closer analysis of the Ub-containing foci showed that while G3-YSD induced ring-shape structures, exposure to C3-YSD generated irregular-shaped foci that are filled with ubiquitinated proteins (Fig. 5E). These observations could suggest that DNA recognition is a first step in the formation of condensates containing ubiquitinated proteins, however, activation of the sensor is required for the assembly of higher-ordered structures. Taken together, our results are consistent with the idea that the ring-shaped structural organization is required for efficient downstream immune signaling.

**Figure 4:**
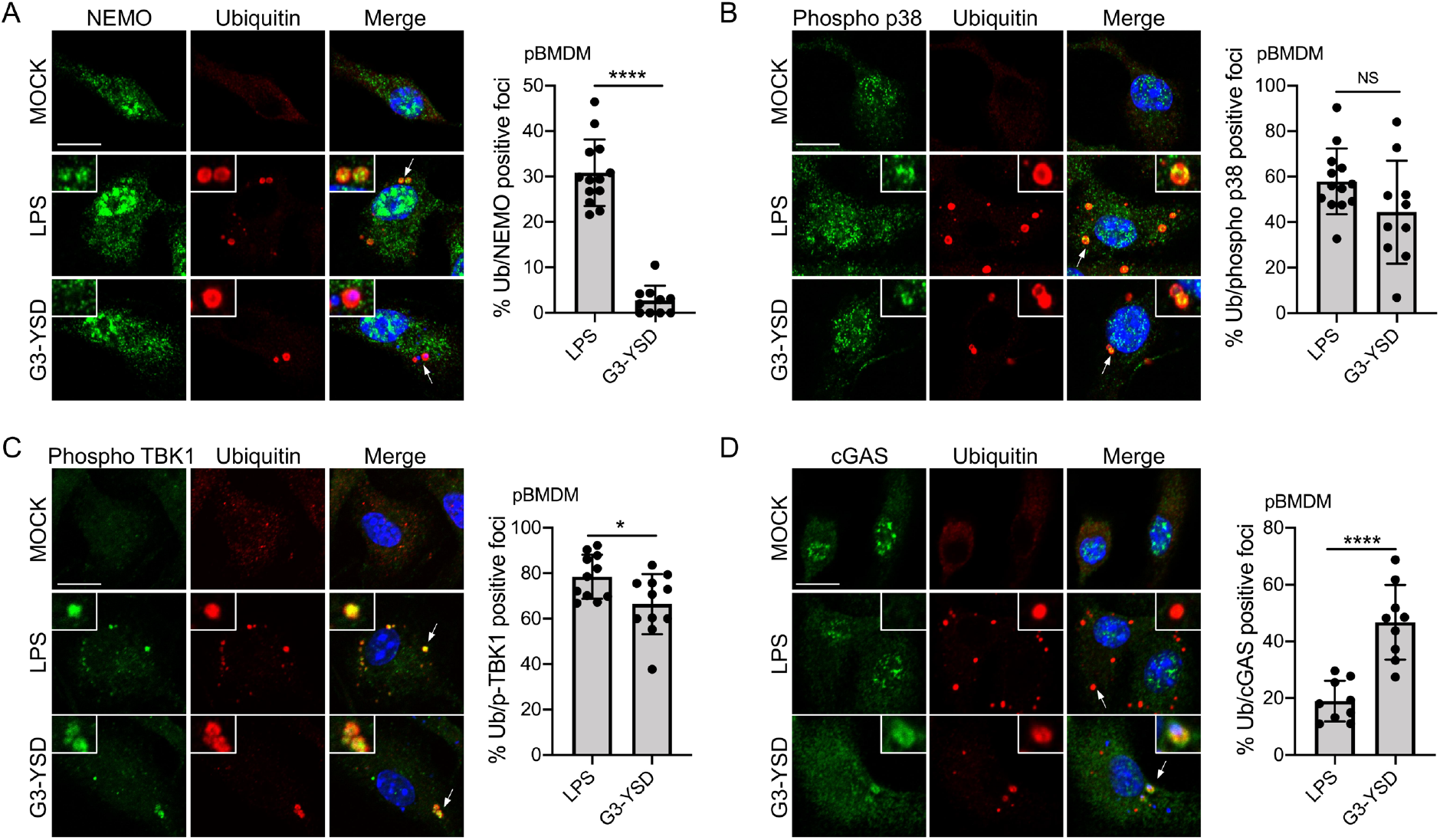
Proteins involved in innate immune pathways are associated with the Ub-containing structures. pBMDM stimulated for 6h with 10 ng/ml of LPS or transfected with 2 μg/ml of G3-YSD were stained using antibodies against polyubiquitinated proteins and NEMO/IKKg (***A***), phosphorylated p38 MAPK (***B***), phosphorylated TBK1 (***C***) or cGAS (***D***). Representative images from three independent experiments are shown. The arrows indicate the regions that are displayed in higher magnification. The quantification was done according to the description in the *materials and methods* and the graphs represent the mean and the corresponding standard deviation of the mean. Significant differences were calculated using two-tailed Student’s t-test (NS, not significant, *p < 0.05, ****p < 0.0001).

**Figure 5:**
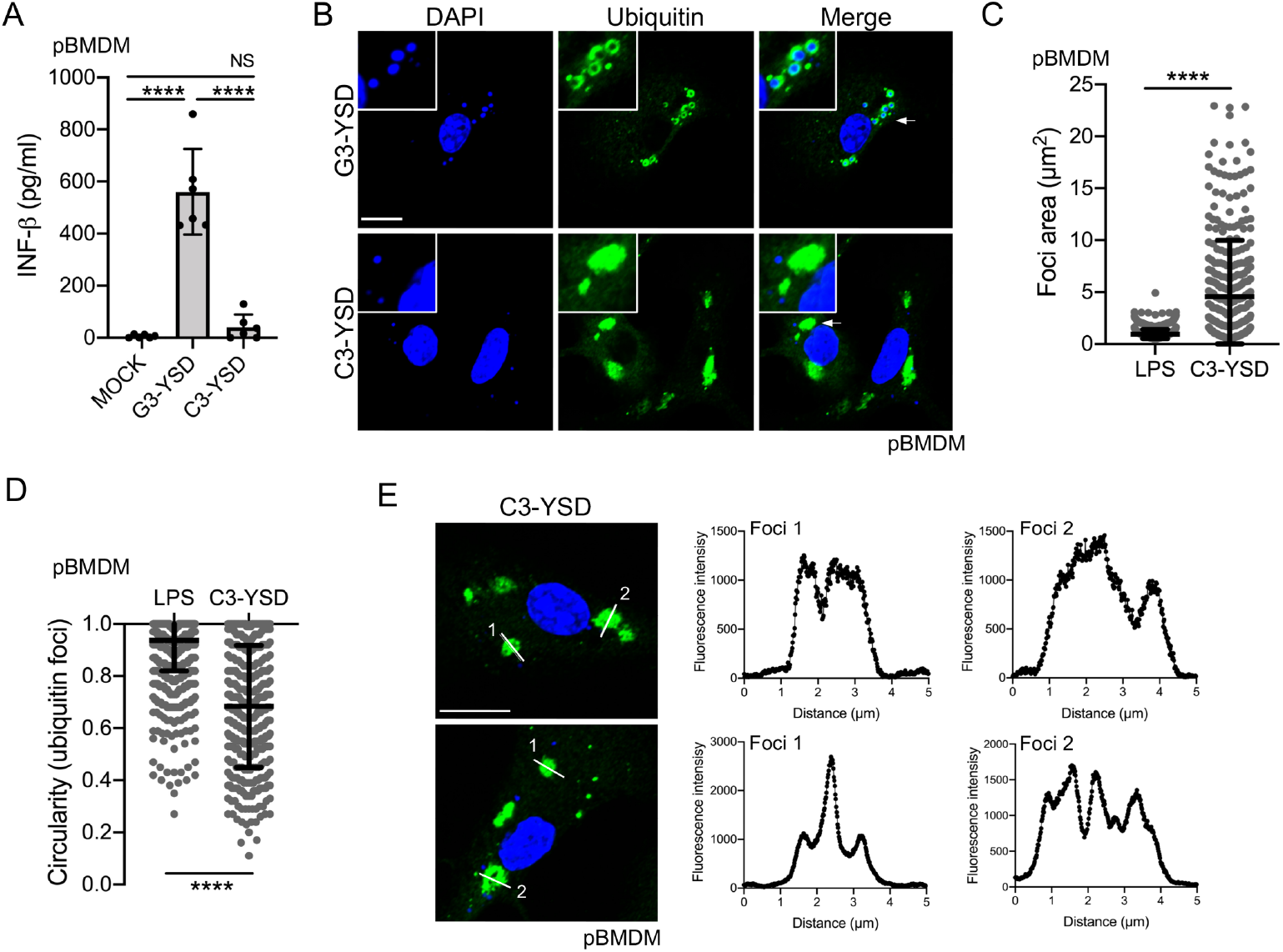
Functional ligand recognition is required for the formation of ubiquitin foci after dsDNA stimulation. ***A***. Quantification of INF-β level in the supernatant of pBMDM transfected for 6h with 2 μg/ml of G3-YSD or C3-YSD. ***B***. Representative images of pBMDM transfected with G3-YSD and C3-YSD 2 μg/ml for 6h and stained for polyubiquitin proteins. The transfected DNA and the nuclei were stained using DAPI. The arrows indicate the region that is displayed in higher magnification. Quantification of the foci area (μm^2^) *(**C**)* and the circularity ***(D)*** using the staining for polyubiquitinated proteins from pBMDM treated with 10 ng/ml of LPS or transfected with 2 μg/ml of C3-YSD for 6h. ***E***. Representative enlarged single plane confocal images showing the structures of foci stained for ubiquitinated proteins after transfection with 2 μg/ml C3-YSD for 6h. The fluorescence intensity was calculated across the white line in the neighboring panel. Significant differences were calculated using one-way ANOVA with Tukey’s post-tests *(A)* or two-tailed Student’s t-test *(C, D)* (NS, not significant, **** p < 0.0001, † significantly lower). Scale bar: 10μm.

## DISCUSSION

Multiple conditions or stresses have been linked to the presence of cytosolic structures that contain ubiquitinated proteins. The best-studied example is the aggresome, which is a large assembly of aggregated and ubiquitinated proteins that are localized at the perinuclear region near the MTOC and occurs primarily when the function of the ubiquitin-proteasome system is altered (Kopito, 2000). Another example is the ALIS foci, which contain ubiquitinated proteins but possess different properties compared to the aggresome, like their size, their cellular localization, and the absence of a vimentin cage surrounding them, among others (Szeto et al., 2006). These condensates were seen in response to multiple stresses, including blocks in protein translation generating truncated proteins unable to fold properly, treatment with heavy metals, exposure to an oxidative environment, and immune stimulation (Jena et al., 2018, Liu et al., 2012, Szeto et al., 2006, Vasconcellos et al., 2016). The current model explaining the formation of these structures, which relies mainly on observations obtained using cellular stressors that have in common to induce protein misfolding, suggests that p62 will recruit free cytosolic polyubiquitinated and unfolded proteins and assemble them into packages. Although the idea of generalizing these findings to all structures containing ubiquitinated proteins independent of the signal triggering their formation is highly attractive, a better understanding of the relationship between the element triggering the formation and the components recruited to these foci is required to fully understand the nature of these structures.

In this study, we used a low concentration of LPS as a physiologic immune trigger to directly study the structural features of these Ub-containing inclusions in the context of immune stimulation. We could observe that foci generated in response to immune stimulation shared structural similarities with the foci generated following proteotoxic stress (Fujita et al., 2011, Jena et al., 2018, Liu et al., 2012, Szeto et al., 2006, Vasconcellos et al., 2016), including the presence of ubiquitinated proteins and Ub-binding proteins like the adaptor protein p62. However, our study provides multiple lines of evidence suggesting that these Ub-containing foci cannot be considered as one-size-fits-all and multiple variations of these structures can be observed depending on the signal triggering their formation. Indeed, although the current model suggests that these foci are an assembly of misfolded proteins, we observed here that immune stimulation of macrophages is not associated with an increase in cytosolic protein aggregation. The presence of aggregated proteins in Ub-containing puncta was described in cells exposed to oxidative stress, which is a condition known to damage proteins and induced aggregation (Jena et al., 2018, Weids et al., 2016). However, macrophages are highly plastic cells that can rapidly adapt their function in response to an environmental challenge (Murray and Wynn, 2011, Wynn et al., 2013). As these cells are a part of the front-line defense against invading microbes, it is not surprising that sensing microbial presence is not associated with the overwhelming of the cellular degradation capacity and formation of micron-size protein aggregates. We also found that immune stimulation resulted in the formation of ring-shaped structures with the ubiquitinated proteins localized on the edges of a structure, an observation at odds with the current model of protein aggregates. Indeed, amorphous protein aggregation described the abnormal and unordered association of misfolded proteins leading to larger aggregates (Hartl et al., 2011). In this scenario, we should observe ubiquitinated proteins all over the inclusions, not only on the edges. The third line of evidence comes from the result that the Nrf2 transcription factor is not essential for the formation of Ub-positive puncta in response to LPS stimulation, in good agreement with the assembly of ALIS foci independently of Nrf2 in response to the autophagy inhibitor 3-MA (Wenger et al., 2012). However, this observation differs from previous studies showing that the Nrf2 signaling pathway was required in response to oxidative stress as well as overnight treatment with a high and non-physiological concentration of LPS (Jena et al., 2018, Vasconcellos et al., 2016, Fujita et al., 2011), suggesting that the requirement for Nrf2 activity is context-dependent. Taken together, these observations are consistent with the idea that distinct signaling pathways can lead to the generation of comparable Ub-containing cytosolic puncta, but possibly serving context-dependent functions.

Although it was previously shown that LPS stimulation and intracellular bacterial infections were sufficient to trigger the formation of Ub-containing foci in macrophages (Liu et al., 2012, Canadien et al., 2005), the extent of signal that can instigate this response in macrophages was still ill-defined. Here, we showed that ligands from both Gram-positive and Gram-negative bacteria, infection with the bacterial pathogens *L. monocytogenes* and MRSA, as well as exposure to cytosolic dsDNA, a hallmark of multiple viral and bacterial infections, are signals for the formation of Ub-containing structures in macrophages. Even though these signals induced the formation of ring-shaped structures containing ubiquitinated proteins surrounding the edges of the foci, major differences could be observed regarding the number of foci per cell, their size, and their circularity. Moreover, the basic core components required for the formation of these structures is dependent on the signal triggering their formation. Indeed, LPS-induced foci are completely dependent on the presence of the adaptor protein p62 whereas Ub-positive foci can be observed in response to cytosolic dsDNA in cells deficient for p62. This observation is somehow reminiscent of the differential role for Nrf2 signaling depending on the trigger and reinforces the idea that the formation of these Ub-containing structures is context-dependent. More importantly, we also observed that a strong agonist is required for the formation of the typical ring-shaped foci following dsDNA sensing, suggesting a direct link between the recognition of the immune stimuli, the formation of the Ub-containing structures, and the resulting immune response. Accordingly, we found a significant enrichment of multiple regulators of major innate immune signaling pathways downstream of either LPS stimulation or cytosolic dsDNA sensing in these Ub-containing structures. This recruitment occurs with a certain degree of specificity as NEMO, a major modulator of the canonical NF-kB pathway, was present only in foci formed downstream of LPS stimulation whereas cGAS was mainly observed in those triggered by dsDNA sensing. Taken together, our results are consistent with the idea that the cytosolic puncta containing ubiquitinated proteins generated after immune stimulation of macrophages are structural platforms containing context-dependent components of major immune signaling pathways.

An emerging concept in innate immunity is the assembly of large oligomeric and modular platforms downstream of microbial detection to regulate host defenses. The SMOCs, which included the myddosome, the triffosome, and the inflammasomes among others, are well-described examples of such complexes. These SMOCs shared the properties of assembling on membranous organelles and often required proteins containing death effector domains (DEDs), pyrin domains (PYRs), or caspase activation and recruitment domains (CARDs) that nucleates helical filament formation by homotypic protein-protein interactions (Kagan et al., 2014, Ha et al., 2020). Although this model for immune signaling is attractive, the SMOCs are not unique in their function of organizing centers for immune responses. Indeed, it was recently shown that multivalent interaction between cGAS and dsDNA induced the formation through phase separation of liquid-like cytosolic foci in which cGAS is concentrated to enhance the production of the second messenger cGAMP required for STING activation and the Type I interferon response (Du and Chen, 2018). Stress granules are another type of membrane-less organelle generated by liquidliquid phase separation that plays the role of organizing platforms for innate immune signaling. Stress granules are dynamic biomolecular condensates that assemble in a context-dependent manner in response to multiple cellular conditions, leading to a variety of functions, from the alteration of mRNA translation and degradation to modulation of signaling pathways and antiviral responses (Alberti et al., 2019, Protter and Parker, 2016). Besides the long-known antiviral role of stress granules in blocking protein translation, multiple antiviral regulatory proteins, including RIG-I, PKR, and RNase L, are recruited to these granules to promote their activation and efficient immune response (Manivannan et al., 2020, Onomoto et al., 2012, Reineke and Lloyd, 2015).

The Ub-containing structures observed here in macrophages might serve a function similar to the ones described for SMOCs or the antiviral stress granules. Ubiquitinated proteins and ubiquitin-binding proteins like p62 might act as a scaffold for the formation of these structures, either by forming oligomeric structures similar to what was observed for the assembly of SMOC complexes or by promoting a liquid-liquid phase separation. As only a few components of these Ub-containing structures are known, it is possible that a major protein, possibly modified through ubiquitination, can assemble into large helical structures required for the formation of micron-size foci in cells. Alternatively, these structures might assemble through phase separation into molecular condensates, where proteins can diffuse freely and retain their native conformation and activities. In agreement with this model, p62 was shown to induce liquid-liquid phase separation in a process that was dependent on the presence of ubiquitin chains (Sun et al., 2018). Undoubtedly, distinguishing between these two models will require further studies and might help shape our understanding of the mechanism behind the assembly of immune signaling complexes in macrophages. An interesting feature of these Ub-positive structures is related to their formation timeline. For comparison, the myddosome assembles minutes after engagement of TLR4 by LPS while these Ub-containing structures started being visible around 4h post-stimulation, suggesting that these two complexes might regulate different stages of the macrophage response to sustain LPS exposure. An alternative model that we cannot exclude to explain our observations is a function for these Ub-containing structures in the modulation of signaling pathway through the sequestration instead of activation of critical components, a suggested function for stress granules in the regulation of the RACK1, TORC1, and TRAF2-dependent signaling pathways (Arimoto et al., 2008, Kim et al., 2005, Takahara and Maeda, 2012). Future work aimed at identifying the components of the Ub-containing structures observed under immune stimulation will be required to better understand their function in the regulation of macrophage immune responses. In summary, our results are consistent with the idea that the cytosolic Ub-containing foci might be subcellular sites for regulation of innate immune signaling in macrophages.

## MATERIALS AND METHODS

### Mice

Mice were housed in specific pathogen-free facilities, maintained by the Unit for Lab Animal Medicine of the University of Michigan. Wild-type C57BL/6 mice (stock No: 000664), Nrf2^-/-^ (stock No:017009), cGAS^-/-^ (stock No: 026554), and Sting^-/-^ (stock No: 025805) were purchased from Jackson Laboratories. The TLR2/4/9^-/-^ femurs were a gift from T. Merkel (FDA) (Hassan et al., 2012). Femurs from male and female WT and p62^-/-^ littermate mice were a gift from J. Moscat (Sanford Burnham Prebys Medical Discovery Institute) (Duran et al., 2004). The Atg7^fl/fl^ and the Atg7^fl/fl^ LysM-Cre femurs were a gift from Dr. J. A. Swanson (University of Michigan Medical School) (Komatsu et al., 2005, Hwang et al., 2012). Whenever possible, independent replicate experiments were done using cells isolated from different mice, including from male and female animals. This study was carried out following the recommendations in the guide for the care and use of laboratory animals of the National Institutes of Health and the protocol was approved by the committee on the care and use of animals of the University of Michigan.

### Primary cell culture and cell lines

Bone-marrow derived macrophages (BMDM) were prepared by flushing mouse femurs in Dulbecco’s modified Eagle’s medium (DMEM) medium. Specific BMDM media containing 50% DMEM, 30% L929-conditioned medium, 20% heat-inactivated fetal bovine serum (FBS), 5% L-Glutamine, 1% sodium pyruvate, and 0.05% β-mercaptoethanol and 100 units/ml of penicillin-streptomycin (Pen/strep) was used for differentiation. Cells were fed fresh media after three days and incubated for an additional three days to complete the differentiation process. To generate immortalized BMDM (iBMDM), bone-marrow cells were transduced with the J2 retrovirus immediately after isolation and differentiated in macrophages as above (Gandino and Varesio, 1990). iBMDM were cultured in DMEM medium supplemented with 10% FBS, 10% L929-conditioned medium, 5% L-Glutamine, 1% sodium pyruvate, 0.05% β-mercaptoethanol and Pen/Strep. Cells were incubated at 37°C with 5% CO_2_.

### Bacterial strains and culture condition

*L. monocytogenes* 10403S was grown in brain heart infusion (BHI) broth statically at 30°C overnight. The community-acquired methicillin-resistant *Staphylococcus aureus* strain USA300 LAC harboring pSarA-GFP (MRSA-GFP) was cultured in tryptic soy medium (TSA, Becton Dickinson) overnight at 37°C with shaking (Boles et al., 2010). Before each experiment, the bacteria were centrifuged, washed once, and diluted in PBS. The OD_600_ was used to determine the inoculum.

### Macrophage infections and treatment

pBMDM were seeded on 6-well plates onto microscope coverslips at a density of 0.5 x 10^6^ cells per well and incubated overnight. The next day, cells were treated in a final volume of 1 ml per well for 6h, unless specified. For immune stimulation, macrophages were treated with 10 ng/ml LPS (InvivoGen, tlrl-smlps), 100 ng/ml Pam3SCK4 (InvivoGen, tlrl-pms) or transfected with 2 μg/ml of G3-YSD (InvivoGen, tlrl-ydna), 2 μg/ml of C3-YSD (InvivoGen, tlrl-ydnac), or 25 pmol of ISD (IDT, sequence: TACAGATCTACTAGTGATCTATGACTGATCTGTACATGATCTACA) using the transfection reagent Turbofect (ThermoFisher, R0531). Alternatively, cells were infected with *L. monocytogenes* at an MOI of 5 for 0.5h. After 3 PBS washes, cells were incubated in fresh media containing 10 μg/ml of gentamycin to kill extracellular bacteria. Similarly, cells were infected with MRSA USA300 at an MOI of 20 for 1h, followed by 3 PBS washes. Fresh media containing 100 μg/ml of gentamycin was added for 20 minutes and replaced by media containing 10 μg/ml gentamycin for the remaining of the infection. For proteasome inhibition, pBMDM were treated for either 6h or 18h with 5 μM MG132 (Cayman Chemicals, S2619). Transcription and translation were blocked by treating the cells for 0.5h with either 1 μg/ml actinomycin D (Sigma-Aldrich, A9415) or 5 μg/ml cycloheximide (Sigma-Aldrich, C7698) before adding LPS at a final concentration of 10 ng/ml. The evaluation of the effect of reactive oxygen species was done by treating the cells for 6h with 10 mM of *N*-Acetyl-L-Cysteine (Millipore, 1009005).

### Immunofluorescence staining and confocal microscopy

After treatment, cells were washed three times with PBS, fixed at room temperature for 20 minutes with 4% paraformaldehyde, and permeabilized with Tris-buffered saline (TBS) with 0.1% Triton X-100 for 10 minutes. All the staining was performed sequentially as strong cross-reactivity was observed, especially when using the guinea pig anti-p62 antibody (ARP, #03-GP62-C). Blocking for 45 minutes was performed at room temperature between each set of primary/secondary antibodies in TBS buffer containing 0.1% Triton X100, 3% bovine serum albumin (BSA), and 10% normal goat serum (ThermoFisher, #100000C). Incubations were done for 1h at room temperature in the staining buffer (TBS with 0.1% Triton X-100 and 3% BSA) for the primary antibodies and 0.5h for the secondary antibodies. Coverslips were extensively washed between each step. The primary antibodies used are the mouse anti-polyubiquitinated conjugates (Enzo Life Sciences, BML-PW8805-0500), the guinea pig anti-p62 (ARP, 03-GP62-C), the rabbit anti-G3BP (Abcam, ab181150), the mouse anti-ubiquitin, Lys48-specific (Millipore, 05-1307), the mouse-anti-ubiquitin, Lys63-specific (Millipore, 05-1308), the rabbit anti-LC3 (MLB International, PM036), the rabbit anti-cGAS (Cell Signaling, 31659), the rabbit anti-IKKg/NEMO (Abcam, ab178872), the rabbit anti-phospho p38 (Thr180/Tyr182) (Cell Signaling, 4511), the rabbit anti-phospho TBK1 (Ser172) (Cell Signaling, 5483) and the rabbit *Listeria* O antisera (ThermoFisher, DF2300). Secondary antibodies used (all from ThermoFisher) are the Alexa Fluor-488 goat anti-mouse IgM (A21042), the Alexa Fluor-594 goat anti-mouse IgM (A21044), the DyLight-650 goat anti-mouse IgM (SA5-101053), the Alexa Fluor-488 goat anti-guinea pig IgG (A11073), the Alexa Fluor-488 goat anti-rabbit IgG (A11034), and the Alexa Fluor-594 goat anti-rabbit IgG (A11037). DAPI was used to stained nucleic acids (ThermoFisher, D1306). Coverslips were mounted on microscope slides using the Prolong Gold mounting reagent (ThermoFisher, P36930). Images were taken on a Nikon A1 confocal microscope using a 60X objective and connected to the Nikon Elements software. The FIJI software was used for image processing (Schindelin et al., 2012). For quantification of the number and the size of the ubiquitin-containing foci, projections of stack confocal images representing 5 μM thick sections of the cells were used. To label the foci, the analyze particle plugin was used with the size of particles set between 0.35 and 30 μm^2^ and the circularity factor above 0.3. Alternatively, the particle analysis was done on all particles with a circularity between 0 and 1 and the shape descriptor plugin was used to evaluate the degree of circularity. For quantification of the structures positive for ubiquitinated proteins and another component (cGAS, phospho-p38, phospho-TBK1, NEMO/IKKg, and LC3), single confocal sections were acquired. The mean fluorescence intensity (MFI) values inside the foci were calculated and compared to the MFI values of the entire cell cytosolic region. Each structure with an MFI 2-fold higher compared to the cytosolic background was considered positive.

### Protein extraction and immunoblotting

pBMDM were seeded on a 6-well plate at a density of 1 x 10^6^ cells per well and incubated overnight. After treatment of pBMDM as described above, whole-cell lysates were prepared using the following lysis buffer: 10 mM Tris-HCl pH8, 150 mM NaCl, 1 % NP-40, 10 mM EDTA pH8, 1 mM DTT, and 1X Roche protease inhibitors. Lysates were incubated on ice for 15 minutes, diluted in 4X sample buffer (Biorad), quickly sonicated, and incubated at 95°C for 10 minutes. Samples were separated by SDS-PAGE and transferred to polyvinylidene fluoride membrane (PVDF, Millipore). Immunoblotting was performed according to the antibody manufacturers’ instructions (cGAS: Cell Signaling, 31659; STING: Proteintech, 19851-1-AP; Actin: Fisher, MS1295P1).

### Statistical analysis

As indicated in figure legends, results represent the mean and the corresponding standard deviation of the mean for at least three independent experiments, unless specified. Statistical analysis was performed using GraphPad Prism 8 software and one-way analysis of variance (ANOVA) with Dunnett or Tukey’s post-tests or unpaired two-tailed student’s t-test were used, as indicated in each legend.

## SUPPLEMENTAL MATERIALS

**Fig. S1** investigates the type of ubiquitin-linkage observed in ALIS and the role of autophagy for ALIS formation. **Fig. S2** describes the role of transcriptional events for ALIS formation. **Fig. S3** examines the structural properties of Ub- and p62-positive foci induced by dsDNA transfection and infection with *L. monocytogenes*. **Fig S4** demonstrates that ALIS formation in response to cytosolic dsDNA is independent of cGAS and STING function.

## ACKNOWLEDGMENTS

This work was supported by National Institutes of Health award 5R33AI102106 (MXO). Research reported in this publication was supported by the National Cancer Institute of the National Institutes of Health under Award Number P30CA046592 by the use of the following Cancer Center Shared Resource(s): Immune Monitoring Core. We also gratefully acknowledge the University of Michigan Microscopy Imaging Laboratory for resources and technical support. We thank all the members of the O’Riordan laboratory at the University of Michigan Medical School for helpful discussions. We thank Dr. T. Merkel (FDA) for providing femurs from TLR2/4/9^-/-^ mice, Dr. J. Moscat (Sanford Burnham Prebys Medical Discovery Institute) for providing cells from *p62*^-/-^ mice, and Dr. J. A. Swanson (University of Michigan Medical School) for cells from *AtgT^fl/fl^* and *Atg7^fl/fl^ LysM-Cre* mice. We also thank Dr. J. A. Swanson (University of Michigan Medical school) for comments on the manuscript.

## AUTHOR CONTRIBUTIONS

M-E.C and V.R. performed experiments. M-E.C and M.X.O designed the experiments, analyzed the results, and wrote the manuscript.

## ABBREVIATIONS

ALIS: aggresome-like induced structures;
dsDNA: double-stranded DNA;
iBMDM: immortalized bone-marrow-derived macrophages;
LPS: lipopolysaccharides;
pBMDM: primary bone-marrow-derived macrophages;
ROS: reactive oxygen species;
SMOC: supramolecular organizing centers;
TLR: Toll-like receptors;
Ub: ubiquitin.

