## Supplemental Figures for "Macrophage aggresome-like induced structures are flexible organizing platforms for immune signaling"

Ubiquitin-positive cytosolic foci in stimulated macrophages are flexible organizing platforms for immune signaling

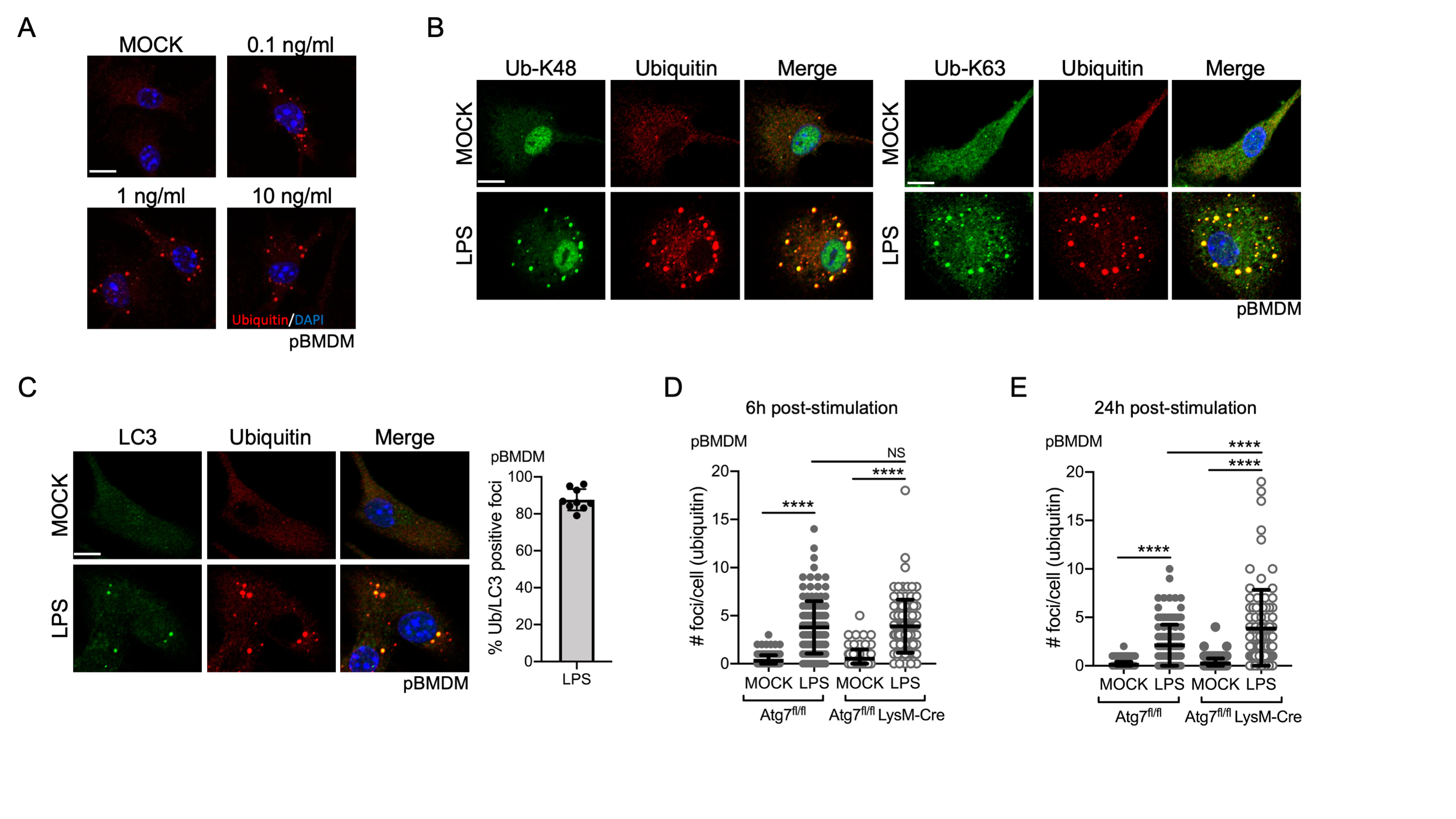

**Figure S1: ALIS foci contain K48- and K63-linked ubiquitin chains and the LC3 protein but their formation is independent of the autophagy machinery. *A***. Representative images of pBMDM treated with 0.1, 1 or 10 ng/ml of LPS and stained for polyubiquitinated proteins. ***B***. Representative images of pBMDM treated with 10 ng/ml of LPS for 6h and stained with antibodies against polyubiquitinated proteins and anti-ubiquitin, Lys48-specific (left panel) or anti-ubiquitin, Lys63-specific (right panel). ***C***. Representative images of pBMDM treated for 6h with 10 ng/ml of LPS and stained with antibodies against polyubiquitinated proteins and LC3. Quantification was done on 10 microscopic fields according to the protocol described in the *methods* section*.* Quantification of the number of foci per cell containing polyubiquitinated proteins from two independent experiments in WT and Atg7^fl/fl^-LysMcre pBMDM treated with LPS 10 ng/ml for 6h (***D***) or 24h (***E***). Graphs represent the mean and the corresponding standard deviation of the mean. Significant differences were calculated using one-way ANOVA with Tukey’s post-test (NS, not significant, ****p < 0.0001). Scale bar: 10µm.

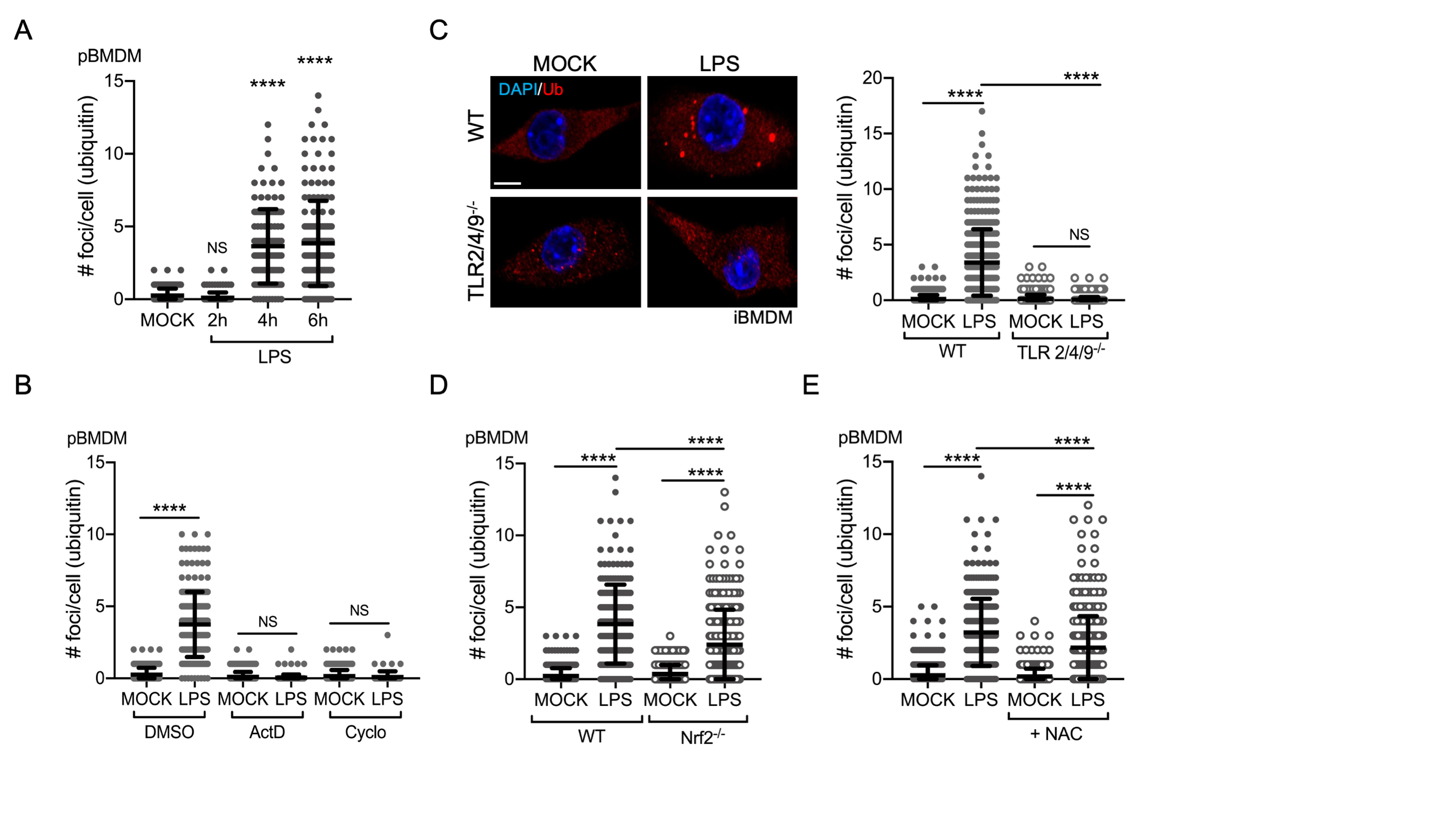

**Figure S2: LPS induces ALIS formation through transcriptional events. *A*.** pBMDM were treated with LPS 10 ng/ml for 2h, 4h or 6h and stained with antibodies against polyubiquitinated proteins. ***B***. Quantification of the number of foci per cell containing polyubiquitinated proteins in pBMDM treated for 0.5h with either 1 µg/ml of actinomycin D or 5 µg/ml of cycloheximide before adding LPS 10 ng/ml for 6h. ***C***. Representative images and quantification of the number of foci per cell for WT and TLR2/4/9^-/-^ iBMDM treated for 6h with 10 ng/ml of LPS and stained with antibodies against polyubiquitinated proteins. ***D***. Quantification of the number of foci per cell containing polyubiquitinated proteins in WT and Nfr2^-/-^ pBMDM left untreated or treated with 10 ng/ml of LPS for 6h. ***E***. Quantification of the number of foci per cell containing polyubiquitinated proteins in pBMDM treated with LPS 10 ng/ml alone or in conjunction with 10 mM of *N*-Acetyl-L-Cysteine. Graphs represent the mean and the corresponding standard deviation of the mean from at least two independent experiments. Significant differences were calculated using two-tailed Student’s t test (*A*) or one-way ANOVA with Tukey’s post-test (*B-E*) (NS, not significant, ****p < 0.0001). Scale bar: 10µm.

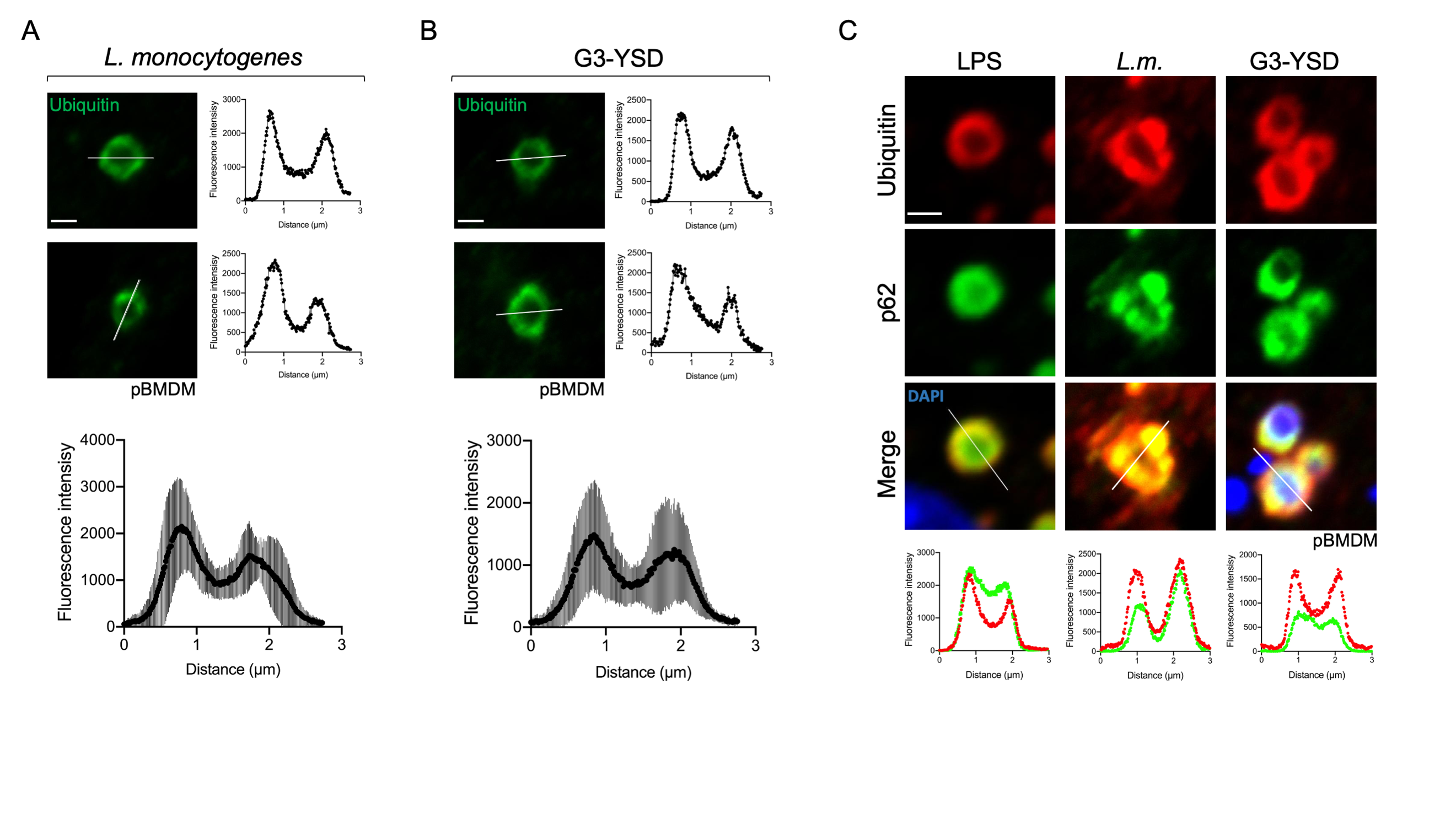

**Figure S3: *L. monocytogenes* infection and exposure to cytosolic dsDNA induce the formation of ring-shaped Ub- and p62-positive structures.** Representative enlarged single confocal images showing the structure of foci stained for polyubiquitinated proteins after infection with *L. monocytogenes* (***A***) or transfection with the dsDNA molecule G3-YSD for 6h (***B***). The fluorescence intensity for the polyubiquitin staining was calculated across the white line in the neighboring panel. The two bottom panels represent the quantification of the polyubiquitin staining fluorescence intensity across 8 individual focus over a distance of 3 µM. The Graphs represent the mean and the corresponding standard deviation of the mean. ***C***. Images representing a magnified area of macrophages stimulated for 6h with 10 ng/ml of LPS, infected with *L. monocytogenes* or transfected with 2 µg/ml of G3-YSD and stained using antibodies against p62 and polyubiquitinated proteins. The fluorescence intensity for the polyubiquitin proteins and p62 staining were calculated across the white line in the panel above.

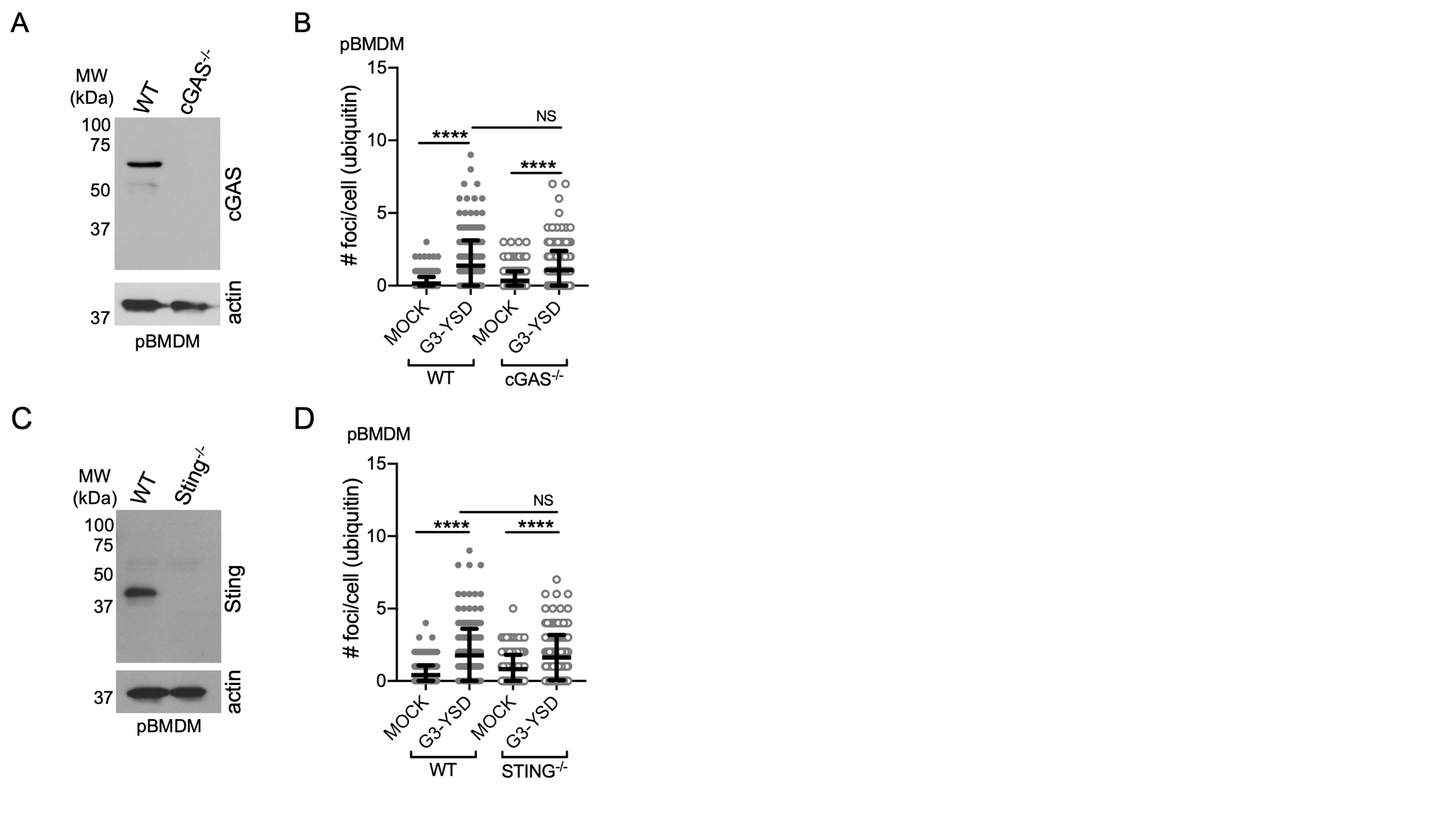

**Figure S4: ALIS formation is independent of cGAS and STING function. *A*.** Immunoblot of WT and cGAS^-/-^ pBMDM whole cell lysates probed with specific antibodies for cGAS and the loading control actin. ***B***. Quantification of the number of foci per cell containing polyubiquitinated proteins in WT and cGAS^-/-^ pBMDM left untreated of transfected for 6h with 2 µg/ml of G3-YSD. ***C***. Immunoblot of WT and STING^-/-^ pBMDM whole cell lysates probed with specific antibodies for STING and the loading control actin. ***D***. Quantification of the number of foci per cell containing polyubiquitinated proteins in WT and STING^-/-^ pBMDM left untreated of transfected for 6h with 2 µg/ml of G3-YSD. Graphs represent the mean and the corresponding standard deviation of the mean from two independent experiments. Significant differences were calculated using one-way ANOVA with Tukey’s post-test (NS, not significant, ****p < 0.0001).
